# Stable isotopes (δ^13^C, δ^15^N, δ^34^S) suggest eelgrass (*Zostera sp.*) foddering of Late Iron Age sheep (*Ovis aries*) in Denmark

**DOI:** 10.64898/2026.04.19.719466

**Authors:** Jonas Holm Jæger, Damon C. Tarrant, Michael P. Richards, Jens Ulriksen, Torben Sarauw, Ole Thirup Kastholm, Julie Nielsen

## Abstract

Stable isotope analysis provides an important tool for reconstructing past livestock management practices and landscape use. However, isotopic data for sheep from Late Iron Age (AD 375/400-1050) Denmark remain limited. Here, we present bulk bone collagen δ¹³C, δ¹⁵N, and δ³⁴S isotope analyses of 27 sheep (*Ovis aries*) from six archaeological sites in Denmark, dated to the Germanic Iron Age (AD 375/400-750) and Viking Age (AD 750-1050).

The analysed sheep exhibit a consistent pattern of enriched δ^13^C values relative to previously published isotopic datasets for Scandinavian livestock, while δ^15^N values display substantial inter-individual variability. Sulfur isotope values fall within moderate ranges consistent with mixed terrestrial and coastal environmental influences. The decoupling of δ^13^C enrichment from elevated δ^15^N values suggests that the observed carbon isotope signal does not reflect marine protein consumption but rather the incorporation of a ^13^C-enriched plant resource into sheep diets.

We propose that eelgrass (*Zostera sp.*), either through direct grazing in coastal environments or supplementary foddering with harvested eelgrass, represents a plausible dietary source to explain this isotopic pattern. The results indicate that Late Iron Age sheep management strategies in Denmark incorporated coastal plant resources within flexible pastoral systems, potentially supporting intensified wool production associated with expanding textile economies.

**Highlights:**

- Stable isotope values of Late Iron Age sheep show some dietary marine input.
- Enriched δ^13^C values suggest eelgrass as supplementary fodder.
- δ^34^S values indicate adaptive grazing across coastal and inland landscapes.

## 1. Introduction

Sheep (*Ovis aries*) have been a common livestock species in Southern Scandinavia since the Early Neolithic (Groß et al., 2024; Sørensen, 2014). During the Bronze Age, new woolly sheep breeds were introduced to Scandinavia, gradually replacing the earlier “primitive” Neolithic sheep breeds (Chessa et al., 2009; Daly et al., 2025; Schroeder et al., 2017). This transition coincided with an increased emphasis on sheep husbandry and the earliest evidence of wool textiles in Denmark (Bender Jørgensen, 1986; Nyegaard, 1996).

The role of sheep appears to have intensified further during the Early Iron Age (500 BC-AD 375/400, when faunal assemblages of domestic livestock at large settlements such as Nørre Tranders (Kveiborg, 2008), Smedegård (B. H. Nielsen et al., 2020), and Nørre Hedegård (Runge, 2009) in Jutland were dominated by sheep. For the Late Iron Age, during the subsequent Germanic Iron Age (AD 375/400-750) and Viking Age (AD 750-1050), sheep husbandry remained central to textile production. The emergence of large settlement complexes with hundreds of pithouses associated with textile craft (Frandsen, 2017; Kastholm, 2025; Sarauw, 2019), an increase in textile tools (Andersson Strand, 2021), and the introduction of the sail (Andersson Strand, 2016; Bender Jørgensen, 2012; Kastholm, 2024, 2025) together suggest that the demand for wool, and more broadly textile fibres, including plant fibres such as flax, had increased substantially.

Sustaining wool production at such levels would have depended not only on flock size, but on management practices that directly affected fibre growth and quality. Nutrition is fundamental for maintaining health, reproductive viability, and productivity in sheep. For example, dietary sulfur plays an important role in wool growth and fibre quality, as sulfur-containing amino acids are critical components of keratin, the protein which makes up wool and hair fibres (De Barbieri et al., 2015; Forcada & Abecia, 2006; Olivier & Olivier, 2006; Qi & Lupton, 1994; Ramos et al., 2019). Consequently, foddering practices or foraging behaviour that increased sulfur intake may have had a tangible impact on sheep flocks where wool production was of economic importance. Therefore, reconstructing foraging behaviour and feeding strategies is essential for understanding how Late Iron Age communities managed their flocks.

Stable isotope analysis provides a means of reconstructing feeding strategies and foraging behaviour in prehistoric livestock husbandry regimes. The combined use of δ^13^C, δ^15^N, and δ^34^S isotope values offers a powerful tool for distinguishing between terrestrial and marine-influenced feeding regimes. While δ^13^C and δ^15^N primarily reflect plant communities and trophic level, δ^34^S values are particularly sensitive to marine inputs from coastal environments. Elevated δ^34^S values in herbivores may therefore signal grazing in coastal ecotones or the use of marine-influenced vegetation as supplementary fodder, including seaweeds or seagrass such as eelgrass (*Zostera sp.*), when interpreted alongside δ^13^C and δ^15^N data.

Stable isotope analysis is a well-established approach for reconstructing past livestock diets and landscape use (Katzenberg, 2008; Makarewicz & Sealy, 2015; Nord & Billström, 2018), and has been widely applied to explore past grazing environments, foddering practices, and flock management strategies (Balasse et al., 2012; Macheridis et al., 2024; Madgwick et al., 2012; Makarewicz, 2014; van der Sluis & Reimer, 2021; Wright, 2014).

While dietary isotope analyses have been applied to investigate animal husbandry practices in Denmark across several periods, including the Early Neolithic (Gron et al., 2024), the Roman Iron Age (Jørkov et al., 2010), and the Viking Age (Swenson, 2019), as well as in studies spanning the Mesolithic to the Viking Age (van der Sluis & Reimer, 2021), isotopic data for sheep dated to the Late Iron Age in Denmark remain limited. At present, published dietary isotope data from this period comprise only a small number of individuals, all dated to the Viking Age, with no isotope data currently available for sheep dated to the Late Iron Age (Swenson, 2019; van der Sluis & Reimer, 2021), severely constraining broader interpretations of sheep husbandry strategies in the latter half of the first millennium AD.

Here, we present the results of bulk bone collagen δ^13^C, δ^15^N, and δ^34^S isotope analyses of 33 individual Late Iron Age sheep from Denmark (Figure 1). All individuals were previously identified to genus by ZooMS (Jæger & Gotfredsen, unpublished), allowing interpretations to be made explicitly within a sheep husbandry framework rather than a potentially mixed ovicaprine dataset.

**Figure 1.**
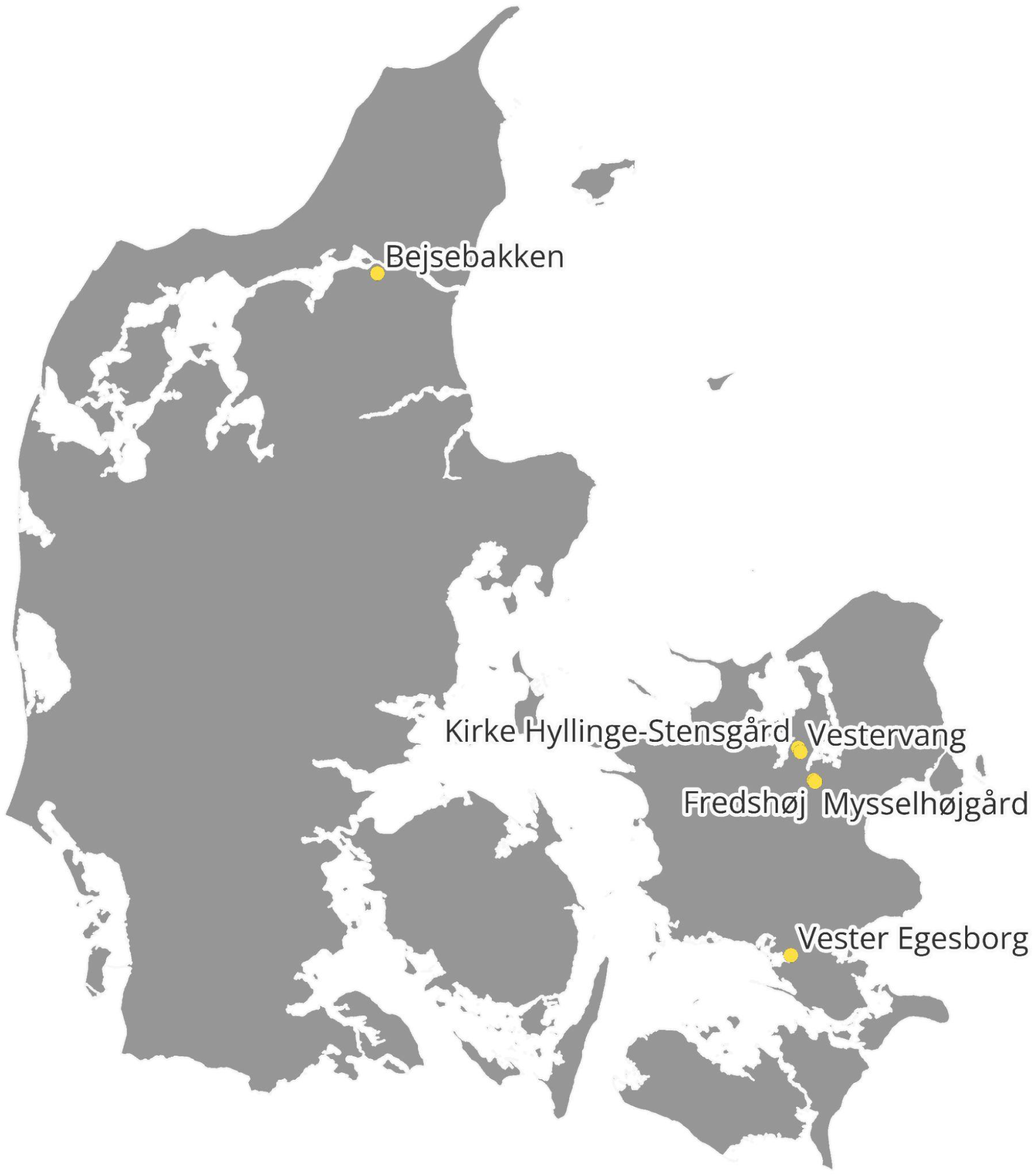
Map of Denmark showing the location of the archaeological sites.

### 1.1. Stable isotope analyses

Carbon (δ^13^C), nitrogen (δ^15^N), and, less frequently, sulfur (δ^34^S) stable isotopes have been widely used in zooarchaeology to reconstruct past sheep husbandry practices, including feeding behaviour, dietary management and landscape use (Balasse et al., 2006; Jørkov et al., 2010; Macheridis et al., 2024; Makarewicz, 2014; Schjerven et al., 2024; Swenson, 2019; van der Sluis & Reimer, 2021; Wright, 2014).

δ^15^N values primarily reflect consumption of animal protein and increase approximately 3-5‰ per increase in trophic level (DeNiro & Epstein, 1981; Depaermentier et al., 2025; O’Connell, 2023; Schoeninger & DeNiro, 1984). However, in herbivores, elevated δ^15^N values may result from several factors, including grazing on manured fields, foddering with vegetation grown on manured soils, the consumption of marine or salt-marsh vegetation, or the trophic enrichment associated with suckling or recent weaning (Balasse & Tresset, 2002; Gron & Rowley-Conwy, 2017; Richards, 2020; Schulting et al., 2017; Schurr, 1998; van der Sluis & Reimer, 2021).

δ^13^C values largely reflect the isotopic composition of consumed vegetation, with only a minor trophic enrichment of approximately 1‰ (DeNiro & Epstein, 1978). In northern European temperate environments, δ^13^C values in herbivores are typically expected to range between -21‰ and -22‰, reflecting a predominantly C3 plant-based diet (Iorga et al., 2021). However, marine and coastal C3 plants are characterised by more enriched δ^13^C values, making carbon isotopes a useful proxy for identifying grazing in coastal or marine-influenced landscapes (Richards, 2020).

δ^34^S isotope values primarily reflect the sulfur isotopic composition of dietary protein, as sulfur is incorporated into bone collagen via methionine, an essential amino acid obtained directly from the diet. As a result, δ^34^S values show minimal trophic fractionation between diet and consumer and closely mirror the isotopic signature of consumed food resources (Nehlich, 2015; Richards et al., 2001).

Marine food webs are characterised by enriched δ^34^S values, typically around 20‰, reflecting the relatively uniform sulfur isotope composition of seawater. Consumption of marine or marine-influenced food resources may therefore result in elevated δ^34^S values relative to terrestrial diets (Nehlich, 2015). However, elevated δ^34^S values in terrestrial herbivores can also be caused by the incorporation of marine-derived sulfur into coastal vegetation via the “sea-spray effect”, i.e. the transport and deposition of aerosolised marine sulfates to terrestrial environments via sea spray, even where diets are entirely terrestrial (Richards et al., 2001; Zazzo et al., 2011). Experimental work on modern sheep has further demonstrated that, under strong sea-spray influence, δ^34^S values may be indistinguishable between animals consuming terrestrial and marine diets, and may show no correlation between δ^13^C or δ^15^N values (E. J. Guiry & Szpak, 2020).

However, in the Southern Scandinavian and Baltic coastal regions, the effect of sea spray is strongly influenced by brackish conditions and substantial freshwater input, resulting in lower and more variable δ^34^S values even in marine fauna (Craig et al., 2006; Göhring et al., 2015; Macheridis et al., 2024). As a result, elevated δ^34^S values cannot be interpreted uncritically as evidence for marine food consumption, such as eelgrass, in this region.

In contrast, low or negative δ^34^S values may indicate dietary inputs derived from freshwater or wetland environments, where anaerobic sulfate reduction produces isotopically depleted sulfur that can be incorporated into plants and transferred to grazing herbivores (E. Guiry et al., 2021; Lamb et al., 2023). When interpreted alongside δ^13^C and δ^15^N, δ^34^S therefore provides valuable contextual information on the environmental setting of dietary resources, rather than a direct or standalone proxy for marine food consumption.

### 1.2. Previous stable isotope analyses of Danish Viking Age sheep

As noted above, only a limited number of stable isotope values have been published for ovicaprines dated to the Viking Age in Denmark (Swenson, 2019; van der Sluis & Reimer, 2021). None of the analysed individuals has been biomolecularly identified to the genus level. While sheep were the dominant ovicaprine species during the Viking Age, and the analysed specimens are therefore likely to represent sheep, the presence of goats (*Capra hircus*) within these datasets cannot be excluded.

The study by van der Sluis and Reimer (2021) includes two ovicaprines from the Limfjord area in Northern Jutland, dated to the Viking Age (AD 800-1050) and the Viking Age/Early Medieval. These individuals exhibit δ^13^C values of -20.5‰ and -21.4‰, and δ^15^N values of 7‰ and 7.1‰, respectively. On this basis, the authors argue that sheep were allowed to roam and graze freely in the surrounding landscape, feeding across different ecotones. The elevated δ^15^N values are interpreted as reflecting the consumption of plants grown on nitrogen-enriched soils, such as manured fields or salt marshes.

Swenson (2019) reports isotope data for two ovicaprine samples from the Viking Age site FHM4573A in Aarhus. The δ^13^C values of these individuals (-21.44‰ and -21.15‰) are broadly consistent with those reported by van der Sluis and Reimer (2021). The δ^15^N values are slightly lower, at 6.86‰ and 6.76‰, respectively; however, δ^15^N values above 6‰ remain consistent with ovicaprines feeding on plants grown on manured soils or salt marsh vegetation (van der Sluis & Reimer, 2021). These results likewise suggest that the ovicaprines from Viking Age Aarhus were managed under similar regimes.

### 1.3. Eelgrass

As discussed above, the isotopic values in Late Iron Age sheep may reflect grazing in coastal environments or the use of supplementary fodder from marine-derived vegetation. In this context, seagrasses such as eelgrass (*Zostera sp*.) represent a particularly relevant resource. Eelgrass is a submerged, flowering plant that forms extensive, dense meadows in shallow coastal waters at depths of up to 20 m below the surface. The coastal waters of modern-day Denmark offer ideal conditions for eelgrass growth due to their relatively low salinity and sheltered nature (McRoy, 1970; Meadows & Fischer, 2024; Thormar et al., 2016).

Besides their ecological importance, eelgrass meadows have historically been an accessible coastal resource. As eelgrass reaches the end of its life cycle, it withers and detaches from the seabed, accumulating along shorelines during autumn and winter, where it can be readily harvested directly from the shore. Alternatively, it can be harvested while submerged (Alm, 2003). Eelgrass has been used extensively for a wide range of purposes in Denmark (J. Nielsen, 2005), the Netherlands (van der Meer, 2009), and Norway (Alm, 2003). Most notably, eelgrass has long been used for thatching. It has also been widely used as stuffing in furniture, mattresses, and pillows, and bedding for livestock, particularly sheep (Alm, 2003). It has long been a well-known fertiliser, and Alm (2003) notes that in the 1700s in Norway, it was considered particularly useful on sandy soils. Importantly, eelgrass has been used as supplementary fodder for livestock, including sheep, which was considered good fodder (Alm, 2003; J. Nielsen, 2005; van der Meer, 2009).

## 2. Materials and methods

### 2.1. Zooarchaeological specimens

Sheep mandibles from six different archaeological sites from the Late Iron Age in Denmark were selected for analysis: Bejsebakken (Sarauw, 2019), Fredshøj (Christensen, 2015, 2024), Kirke Hyllinge-Stensgård (Ulriksen, 1998), Mysselhøjgård (Christensen, 2015, 2024), Vester Egesborg (Ulriksen, 2018) and Vestervang (Kastholm, 2009, 2012). The selected sites represent a range of settlement types and environmental contexts, including rural settlements and sites associated with central place complexes, as well as locations at varying distances from the coast, enabling the investigation of how landscape use and access to coastal resources may have influenced sheep management strategies. All mandibles had previously been identified to genus using ZooMS (Jæger & Gotfredsen, unpublished), ensuring taxonomic reliability, enabling the integration of isotope and palaeoproteomic datasets. The analysed specimens are curated at the Natural History Museum of Denmark, University of Copenhagen, with the exception of the material from Bejsebakken, which is housed at Nordjyske Museer, Aalborg, Denmark.

Specimens were selected from the available biomolecularly identified sheep mandibles, with preference given to mandibles exhibiting good macroscopic preservation and minimal taphonomic alteration. Uneven sample sizes across sites reflect differences in assemblage composition, preservation and the availability of ZooMS-identified mandibles. Contextual information for each specimen is provided in Supplementary Table S1.

### 2.2. Coastal proximity

To provide an environmental context for potential marine influences on isotopic values, the distance between each site and the present-day coastline was calculated using QGIS (v. 3.44.4-Solothurn). Site locations were defined using published coordinates and archaeological documentation, and distances were measured as the shortest linear distance between each site and the nearest coastline feature. Distance to the coast was treated as a contextual environmental variable reflecting potential accessibility to coastal landscapes rather than a direct proxy for grazing areas, as sheep may have been seasonally moved between grazing areas or provided with transported fodder resources.

It should be noted that relative sea-level change and local geomorphological processes may have altered shoreline positions since the Late Iron Age. In Denmark, however, such changes are generally limited, with only minor localised variation. In the Roskilde Fjord in particular, sea-level differences between the Late Iron Age and the present are estimated to be minimal at approximately 0-0.5 m (Kastholm, 2024). Measured distances can therefore be regarded as reasonable approximations of coastal proximity (Table 1).

**Table 1:**
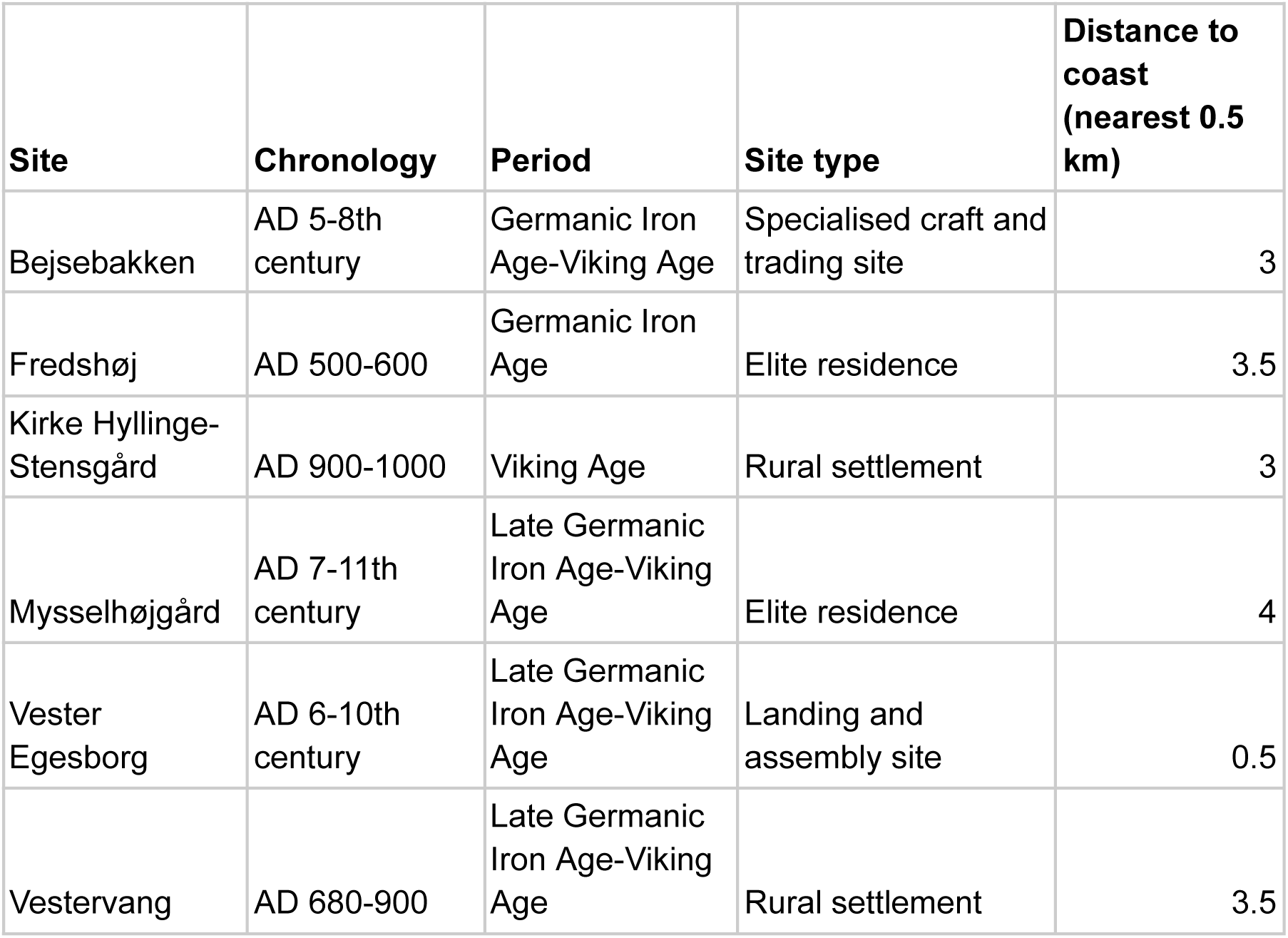
Archaeological sites included in the study with relative chronological attribution and distance to the nearest coastline. Distances were calculated in QGIS and rounded to account for spatial and palaeogeographic uncertainty.

| Site | Chronology | Period | Site type | Distance to coast (nearest 0.5 km) |
| --- | --- | --- | --- | --- |
| Bejsebakken | AD 5-8th century | Germanic Iron Age-Viking Age | Specialised craft and trading site | 3 |
| Fredshøj | AD 500-600 | Germanic Iron Age | Elite residence | 3.5 |
| Kirke Hyllinge-Stensgård | AD 900-1000 | Viking Age | Rural settlement | 3 |
| Mysselhøjgård | AD 7-11th century | Late Germanic Iron Age-Viking Age | Elite residence | 4 |
| Vester Egesborg | AD 6-10th century | Late Germanic Iron Age-Viking Age | Landing and assembly site | 0.5 |
| Vestervang | AD 680-900 | Late Germanic Iron Age-Viking Age | Rural settlement | 3.5 |

### 2.3. Collagen extraction

Collagen was extracted following the modified Longin method (Brown et al., 1988; Longin, 1971). Samples were cut into small pieces weighing approximately ∼200mg, and had the surfaces abraded to remove surficial contaminants. Samples were then demineralised in 0.5 mol HCl at 4°C. Once demineralised, the samples were rinsed three times with MilliQ ultra pure H_2_O. Samples were then solubilised in pH 3 HCl at 72°C for 48 hours. Samples were then filtered using Eeze filters and 30 kDa filters. Collagen greater than 30 kDa was used for isotopic analysis. Collagen aliquots were then frozen and lyophilised for 72 hours.

### 2.4. Isotope ratio mass spectrometry

Collagen was weighed to 1.45mg in tin capsules, and measured via continuous flow stable isotope ratio mass spectrometry in the Isotope Laboratory in the Department of Archaeology at Simon Fraser University. δ^15^N, δ^13^C, and δ^34^S were measured simultaneously on a Thermo Fisher Delta V Plus (Sayle et al., 2019). Samples were normalised to international scales using USGS40 and USGS41 for δ^15^N (AIR) and δ^13^C (VPDB), and S-1 and NBS-127 for δ^34^S (VCDT). Internal lab standards were measured throughout the run SRM3 (Bovine Collagen: δ^15^N: 9.12‰, δ^13^C: -15.55‰, and δ^34^S: 6.3‰) and Casein (δ^15^N: 5.85‰, δ^13^C: -27.26‰, and δ^34^S: 7.56‰): average analytical precision through the run was δ^13^C: 0.44, δ^15^N: 0.01‰, δ^34^S 0.24‰, our reproducibility of SRM measurement was δ^15^N: 0.06‰, δ^13^C: 0.21‰, and δ^34^S: 0.34‰.

## 3. Results

### 3.1. Success ratio

33 samples were selected for stable isotope analysis. Of these, 27 yielded paired δ^13^C and δ^15^N measurements, and 26 yielded δ^34^S measurements. Six samples were excluded from further analysis due to poor collagen preservation and/or unacceptable elemental ratios.

For the samples included in the analysis, all C:N ratios fell within the accepted range of 2.9-3.6 for well-preserved mammalian bone collagen (Schoeninger & DeNiro, 1984). Similarly, all samples analysed for sulfur isotope composition displayed C:S ratios within the accepted range of 600 ± 300 for mammalian bone collagen (Nehlich & Richards, 2009). The results of the stable isotope analysis are summarised in Table 2:

**Table 2.** Sample information and stable carbon, nitrogen and sulfur compositions of sheep mandibles from Bejsebakken, Fredshøj, Kirke Hyllinge-Stensgård, Vester Egesborg and Vestervang. ZMK no. refers to the collection number assigned by the Natural History Museum of Denmark, University of Copenhagen. Additional contextual information is provided in Supplementary Table S1.

| Sample | Site | ZMK no. | Species | Element | % Collagen<br>>30kD | δ13C | δ15N | δ34S | wt%<br>C | wt%<br>N | %S | C:<br>N | C:S |
| --- | --- | --- | --- | --- | --- | --- | --- | --- | --- | --- | --- | --- | --- |
| i008 | Vester Egesborg | ZMK<br>35/2009 | *Ovis*<br>*aries* | Mandible | 10.7 | -20.1 | 6 | 10.6 | 37.1 | 14.4 | 0.3 | 3 | 419.6 |
| i022 | Vester Egesborg | ZMK<br>35/2009 | *Ovis*<br>*aries* | Mandible | 10 | -20.8 | 9.6 | 10.7 | 36.9 | 13.9 | 0.2 | 3.1 | 426.1 |
| i023 | Vester Egesborg | ZMK<br>35/2009 | *Ovis*<br>*aries* | Mandible | 11.9 | -19.7 | 6.1 | 9.6 | 36.6 | 13.7 | 0.3 | 3.1 | 427.6 |
| i200 | Kirke Hyllinge-Stensgård | ZMK<br>80/2003 | *Ovis*<br>*aries* | Mandible | 14.2 | -22 | 6.2 | 9.3 | 36.3 | 11.9 | 0.2 | 3.6 | 401.4 |
| i212 | Kirke Hyllinge-Stensgård | ZMK<br>80/2003 | *Ovis*<br>*aries* | Mandible | 2.1 | -21.7 | 6.7 | 10.1 | 20.7 | 7.7 | 0.2 | 3.1 | 289.3 |
| i315 | Vestervang | ZMK<br>17/2008 | *Ovis*<br>*aries* | Mandible | 11.5 | -20.6 | 6.2 | 11.2 | 39.1 | 14.9 | 0.2 | 3.1 | 471.4 |
| i500 | Fredshøj | ZMK<br>20/2009 | *Ovis*<br>*aries* | Mandible | 1.4 | -20.4 | 6.3 | 10 | 32 | 12.4 | 0.2 | 3 | 373.9 |
| i501 | Fredshøj | ZMK<br>20/2009 | *Ovis*<br>*aries* | Mandible | 5.7 | -19.3 | 5.8 | 9.3 | 33.9 | 13 | 0.3 | 3 | 338.4 |
| i502 | Fredshøj | ZMK<br>20/2009 | *Ovis*<br>*aries* | Mandible | 8.5 | -21.2 | 6.3 | 10.4 | 38.6 | 14.5 | 0.3 | 3.1 | 390.3 |
| i509 | Fredshøj | ZMK<br>20/2009 | *Ovis*<br>*aries* | Mandible | 6.8 | -20.3 | 4.6 | 10.9 | 35.6 | 14.2 | 0.3 | 2.9 | 340.1 |
| i511 | Fredshøj | ZMK<br>20/2009 | *Ovis*<br>*aries* | Mandible | 7.5 | -20.3 | 9.4 | 10.1 | 39.9 | 15.6 | 0.3 | 3 | 413.9 |
| i519 | Fredshøj | ZMK<br>20/2009 | *Ovis*<br>*aries* | Mandible | 4.8 | -20.2 | 6.9 | 9.6 | 36.4 | 14.7 | 0.3 | 2.9 | 374.4 |
| i520 | Fredshøj | ZMK<br>20/2009 | *Ovis*<br>*aries* | Mandible | 1.6 | -19.6 | 6.3 | 10.4 | 26.1 | 10.4 | 0.2 | 2.9 | 320.5 |
| i523 | Fredshøj | ZMK<br>20/2009 | *Ovis*<br>*aries* | Mandible | 8.3 | -20.5 | 6.7 | 10.6 | 40 | 15.5 | 0.3 | 3 | 411.8 |
| i524 | Fredshøj | ZMK<br>20/2009 | *Ovis*<br>*aries* | Mandible | 8.6 | -19.2 | 6.3 | 8.1 | 38.4 | 14.9 | 0.4 | 3 | 313.9 |
| i526 | Fredshøj | ZMK<br>20/2009 | *Ovis*<br>*aries* | Mandible | 7 | -19.2 | 6.4 | 8.9 | 37.3 | 14.6 | 0.3 | 3 | 365.8 |
| i529 | Fredshøj | ZMK<br>20/2009 | *Ovis*<br>*aries* | Mandible | 11.7 | - | - | 10.3 | 38.9 | 11.3 | 0.2 | 4 | 486.5 |
| i606 | Myssehøjgård | ZMK<br>88/1989 | *Ovis*<br>*aries* | Mandible | 5.6 | -20.6 | 6.6 | 9.9 | 37.3 | 14 | 0.2 | 3.1 | 449.6 |
| i618 | Myssehøjgård | ZMK<br>88/1989 | *Ovis*<br>*aries* | Mandible | 11.6 | -19.7 | 7.7 | 9.9 | 38 | 14.8 | 0.2 | 3 | 457.3 |
| i800 | Bejsebakken | ÅHM3984 | *Ovis*<br>*aries* | Mandible | 7.4 | -20.6 | 6.8 | 11.1 | 32.2 | 12.6 | 0.2 | 3 | 471.6 |
| i801 | Bejsebakken | ÅHM3984 | *Ovis*<br>*aries* | Mandible | 6.8 | -19.3 | 5.9 | 11 | 31.5 | 12.4 | 0.2 | 3 | 531.9 |
| i804 | Bejsebakken | ÅHM3984 | *Ovis*<br>*aries* | Mandible | 5.9 | -17.1 | 6.7 | 11.8 | 32.2 | 12.8 | 0.2 | 2.9 | 470.6 |
| i805 | Bejsebakken | ÅHM3984 | *Ovis*<br>*aries* | Mandible | 4.8 | -20.8 | 9.3 | - | 31.3 | 11.8 | 0.4 | 3.1 | 226.8 |
| i807 | Bejsebakken | ÅHM3984 | *Ovis*<br>*aries* | Mandible | 10 | -19.3 | 4.8 | 12.6 | 36.2 | 14.2 | 0.2 | 3 | 525.2 |
| i809 | Bejsebakken | ÅHM3984 | *Ovis aries* | Mandible | 2.3 | -18.8 | 4.9 | 10.6 | 27.9 | 11 | 0.2 | 2.9 | 463.8 |
| i843 | Bejsebakken | ÅHM3984 | *Ovis aries* | Mandible | 9.5 | -20.5 | 6.8 | - | 35.8 | 13.8 | 0.4 | 3 | 217.6 |
| i855 | Bejsebakken | ÅHM3984 | *Ovis aries* | Mandible | 4.8 | -20.2 | 4.8 | 9.6 | 34.9 | 13.2 | 0.3 | 3.1 | 317 |

### 3.2. δ^13^C and δ^15^N compositions

The sheep analysed in this study exhibit a broad range of δ^13^C and δ^15^N stable isotope values (Figure 2), although the degree of variability differs markedly between the two isotope systems. δ^13^C values range from -22.0 to -17.1‰, with a mean of -20.1‰ (± 0.1‰ SD), while δ^15^N values range from 4.6 to 9.6‰, with a mean of 6.5‰ (± 1.3‰ SD).

**Figure 2.**
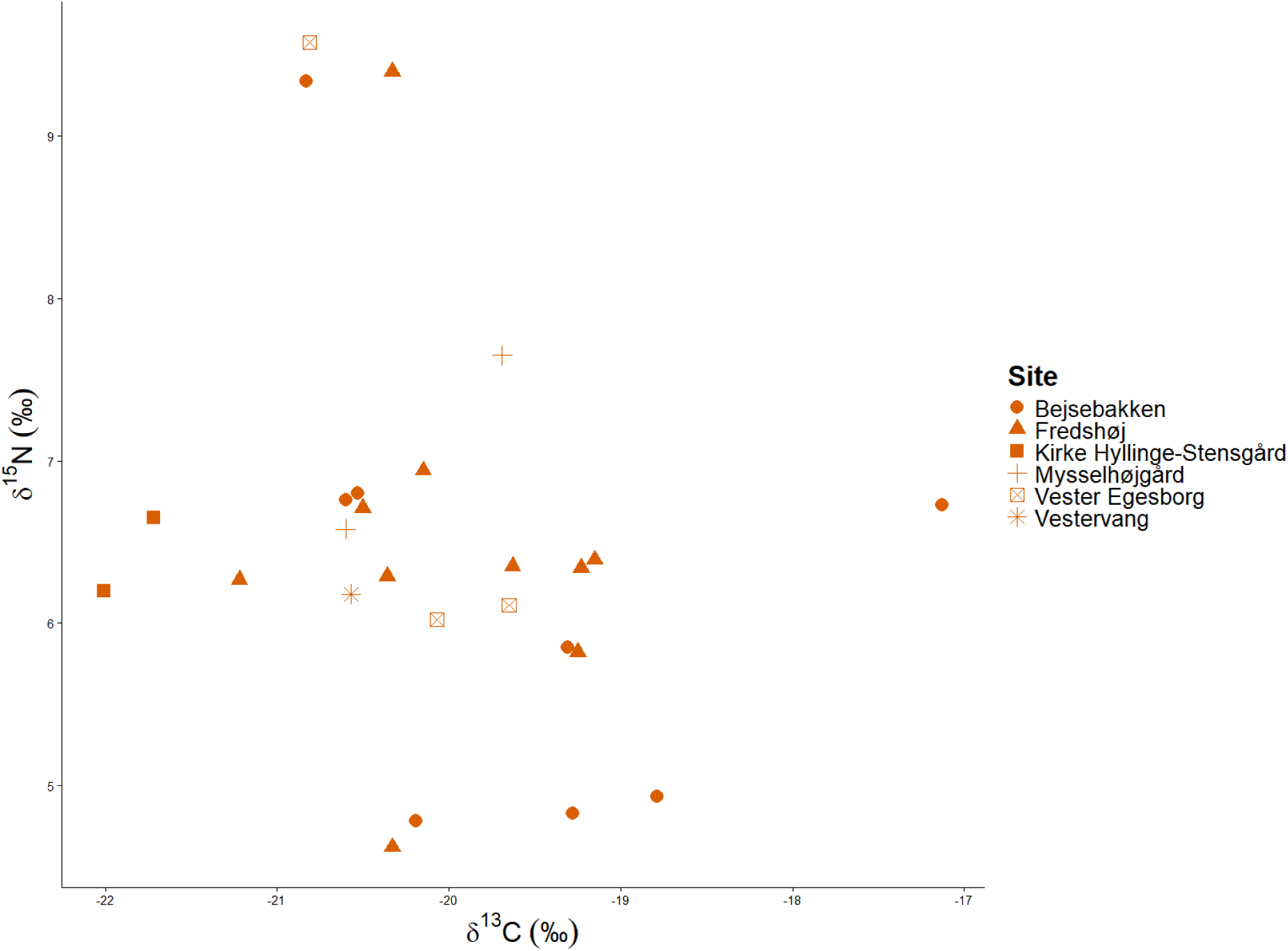
Plot of δ13C and δ^15^N values for all samples analysed for this study.

The δ^13^C values of the analysed sheep display a broad overall range, spanning from -22 to -17.1‰ with a mean of -20.1‰ (± 0.1‰ SD). Most individuals cluster between approximately -20.6 and -19.2‰, indicating a constrained carbon isotope signal across the assemblage. While several individuals exhibit δ^13^C values comparable to previously published values for Early and Late Iron Age domesticated livestock in Denmark (-22.5 to -20.5‰) (Jørkov et al., 2010; Swenson, 2019; van der Sluis & Reimer, 2021), the majority of the analysed sheep show enriched δ^13^C values of -20.5‰ or higher. 14 individuals display enriched δ^13^C values between -20.2 and -17.1‰, exceeding the upper range previously reported for Danish Iron livestock. One individual exhibits a particularly enriched δ^13^C (-17.1‰); however, the overall tight clustering of δ^13^C values indicates that elevated δ^13^C is a consistent pattern rather than a small number of anomalous samples.

In contrast, the δ^15^N values exhibit a greater dispersion than the δ^13^C values, ranging from 4.6 to 9.6‰ with a mean of 6.5‰ (± 1.3‰ SD) (Figure 2). Both low and high δ^15^N values are represented across the assemblage, with several individuals clustering near the lower end of the range (approximately 4.6 to 5.0‰) and others displaying markedly elevated values approaching 9.5‰. In contrast to the constrained δ^13^C values, δ^15^N values show pronounced inter-individual variability. Nevertheless, the overall range of δ^15^N values falls within those previously published for Danish Early and Late Iron Age domestic livestock (4.2-10.7‰) (Jørkov et al., 2010; Swenson, 2019; van der Sluis & Reimer, 2021)

### 3.3. δ^34^S compositions

The δ^34^S values of the analysed sheep also show moderate variability, ranging from 8.1 to 12.6‰, with a mean of 10.2‰ (SD ± 0.9‰) (Figure 3-4). Most individuals cluster between 9.0 and 11.2‰, while one individual exhibits a markedly elevated δ^34^S value of 12.6‰. Although δ^34^S values show greater dispersion than δ^13^C values, they are less variable than δ^15^N. Although a small number of samples lack δ^34^S measurements, the observed range and clustering indicate a generally coherent sulfur isotope signal across the assemblage.

**Figure 3.**
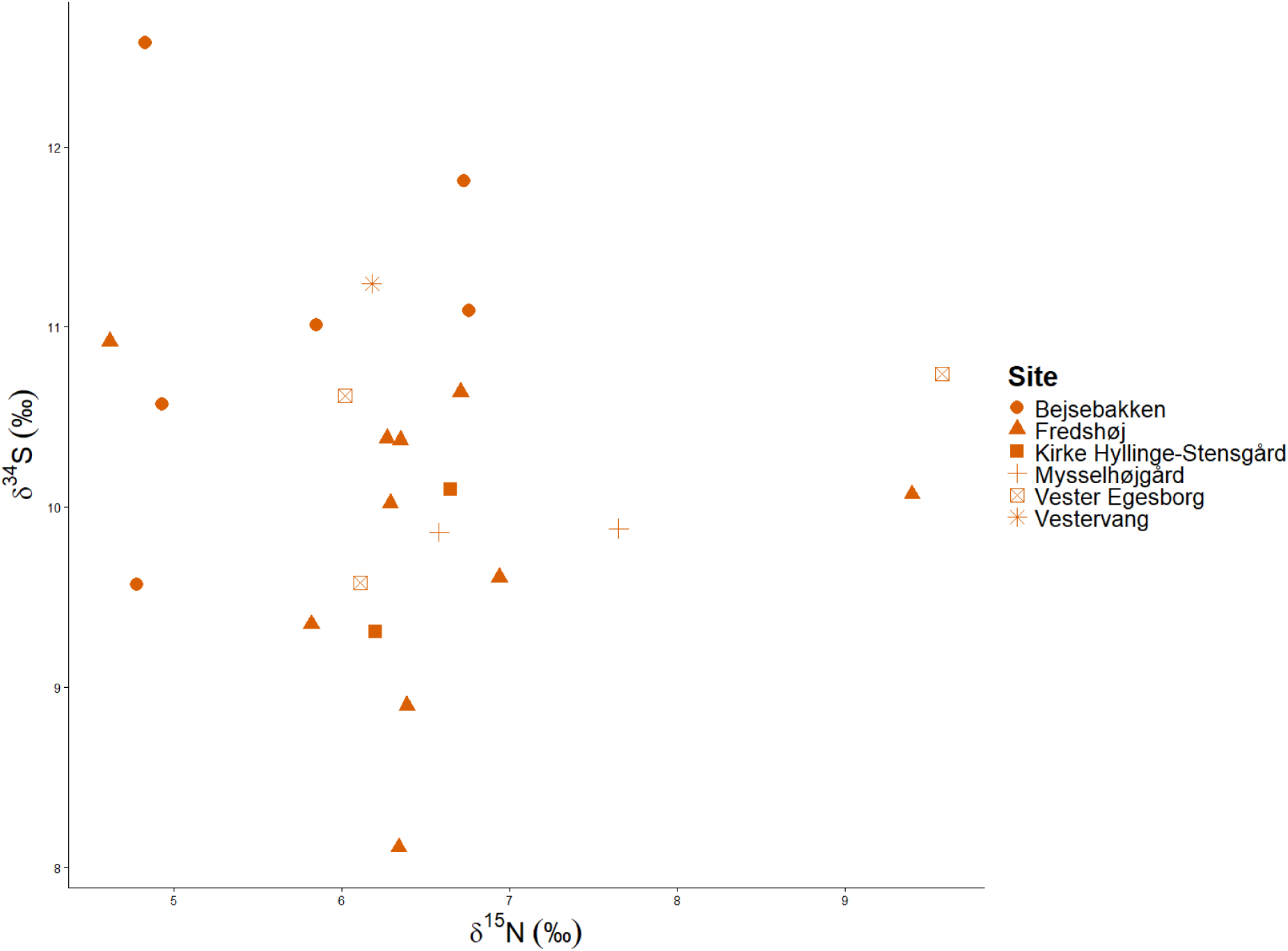
Plot of δ^15^N and δ^34^S values for all samples analysed for this study.

**Figure 4.**
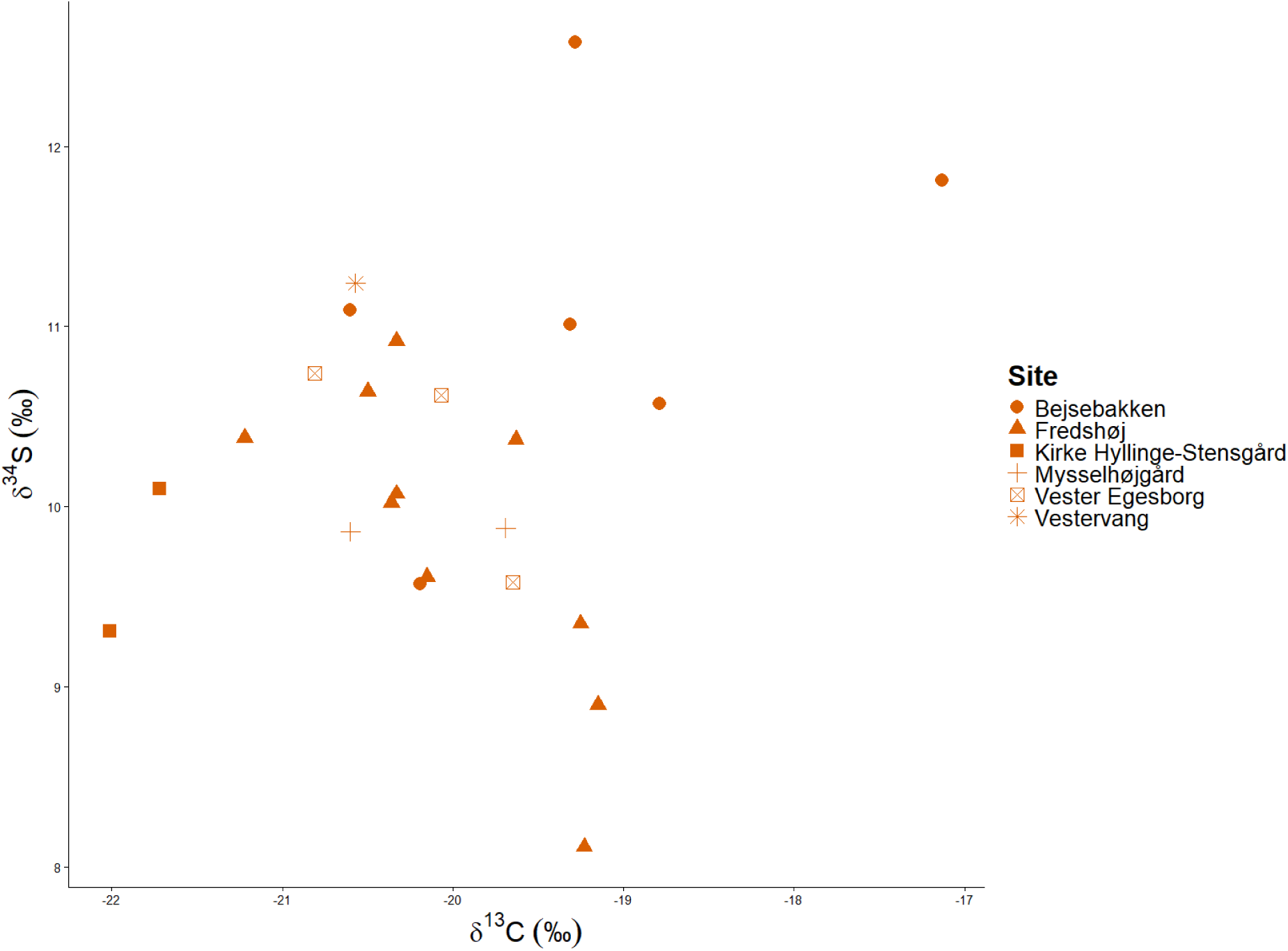
Plot of δ^13^C and δ^34^S values for all samples analysed for this study.

### 3.4. Within-site comparison

Within-site comparisons must be considered in light of uneven sample sizes across sites: Fredshøj and Bejsebakken are represented by comparatively larger datasets, while Mysselhøjgård and Kirke Hyllinge-Stensgård are represented by only two individuals each, and Vestervang is represented by a single individual. Consequently, patterns observed at sites with small sample sizes should be regarded as indicative rather than representative of broader site-level variability.

3.4.1. *Bejsebakken*

At Bejsebakken (*n* = 8), δ¹³C values range from -20.8 to -17.1‰ with a mean of -19.6‰ (± 1.2‰ SD), representing the widest δ¹³C range observed within a single site. δ¹⁵N values span from 4.8 to 9.3‰ with a mean of 6.3‰ (± 1.5‰ SD), indicating substantial inter-individual variability. δ³⁴S values range from 9.6 to 12.6‰ with a mean of 11.1‰ (± 1.0‰ SD), including the most enriched δ³⁴S value recorded in the dataset.

3.4.2. *Fredshøj*

At Fredshøj (*n* = 12), twelve samples yielded δ³⁴S values, while eleven samples produced reliable δ¹³C and δ¹⁵N measurements. δ¹³C values range from -21.2 to -19.2‰ with a mean of -20.0‰ (± 0.7‰ SD), indicating moderate variability relative to Bejsebakken. δ¹⁵N values range from 4.6 to 9.4‰, with a mean of 6.5‰ (± 1.2‰ SD), encompassing both low and high values across the dataset. δ³⁴S values range from 8.1 to 10.9‰ with a mean of 9.9‰ (± 0.8‰ SD), showing moderate dispersion.

3.4.3. *Kirke Hyllinge-Stensgård*

From Kirke Hyllinge-Stensgård (*n* = 2), both samples yielded sufficient collagen for isotopic measurements. The samples yielded δ^13^C values from -22.0 to -21.7‰ with a mean of -21.9‰ (± 0.2‰ SD). δ^15^N values ranged from 6.2 to 6.7‰ with a mean of 6.4‰ (± 0.3‰ SD), and δ^34^S values of 9.3 and 10.1‰ with a mean of 9.7‰ (± 0.6‰ SD). Both individuals fall within the broader ranges observed across the assemblages, and the narrow intra-site variability suggests broadly similar dietary and/or grazing conditions for both individuals.

3.4.4. *Mysselhøjgård*

At Mysselhøjgård (*n* = 2), δ¹³C values range from -20.6 to -19.7‰ with a mean of -20.1‰ (± 0.7‰ SD). δ¹⁵N values range from 6.6 to 7.7‰ with a mean of 7.1‰ (± 0.8‰ SD), indicating limited variability. Both samples yielded identical δ³⁴S values of 9.9‰, suggesting a highly consistent sulfur isotope signal at this site.

3.4.5. *Vester Egesborg*

At Vester Egesborg (*n* = 3), δ¹³C values range from -20.8 to -19.7‰ with a mean of -20.2‰ (± 0.6‰ SD). δ¹⁵N values range from 6.0 to 9.6‰ with a mean of 7.2‰ (± 2.0‰ SD), indicating marked inter-individual variability. δ³⁴S values range from 9.6 to 10.7‰ with a mean of 10.3‰ (± 0.6‰ SD).

3.4.6. *Vestervang*

The single analysed specimen from Vestervang (*n* = 1) yielded δ¹³C, δ¹⁵N, and δ³⁴S values of -20.6‰, 6.2‰, and 11.2‰, respectively, all of which fall within the ranges observed at other sites in the study.

## 4. Discussion and conclusions

The consistency and degree of enriched δ^13^C values in the analysed sheep stand in contrast to previously reported δ^13^C values in livestock from Denmark (Jørkov et al., 2010; Swenson, 2019; van der Sluis & Reimer, 2021). This pattern is evident at the assemblage level and at sites represented by larger sample sizes, indicating that the apparently consistent carbon isotope enrichment across the dataset is unlikely to be driven by a small number of anomalous individuals. Rather, it suggests that δ^13^C enrichment was a systematic component of sheep management practices in Denmark during the Late Iron Age.

### 4.1. Carbon isotope enrichment

Despite spanning a relatively broad absolute range, δ^13^C values are clustered across the dataset, suggesting that enriched carbon isotope values reflect a shared aspect of sheep management practices rather than sporadic or site-specific behaviours. The degree of enrichment observed exceeds the upper range previously reported for contemporaneous domestic livestock, indicating that the sheep analysed for this study were subject to environmental or dietary conditions that differed consistently from those documented in earlier studies (Figure 5) (Jørkov et al., 2010; Swenson, 2019; van der Sluis & Reimer, 2021).

**Figure 5.**
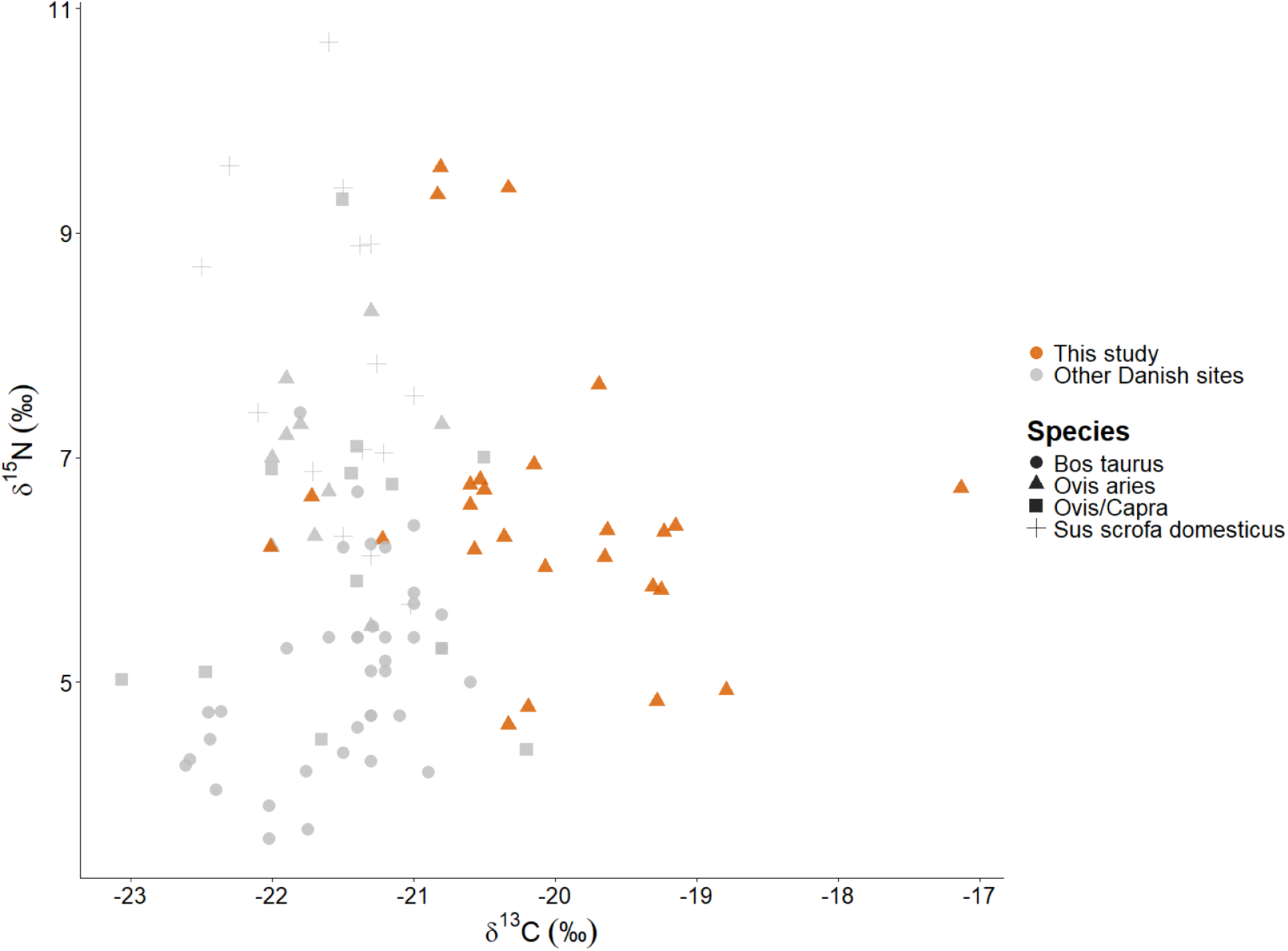
Plot of δ^13^C and δ^15^N values for all samples analysed for this study compared to domestic livestock dated from the Early Iron Age to the Viking Age from previously published studies (Jørkov et al., 2010; Swenson, 2019; van der Sluis & Reimer, 2021).

The δ^13^C values observed in this study (-22.0 to -17.1‰) span both the most negative and most enriched values currently reported for Late Iron Age ovicaprines in Denmark. While similarly depleted values have been documented in Early Neolithic ovicaprines from Syltholm II, where δ¹³C values reach -24.47‰ (Gron et al., 2024), the enriched values recorded here exceed those previously reported for Danish prehistoric ovicaprines. Comparable enrichment has been documented in modern sheep from Orkney, where sheep subsisting on seaweed display a range of δ¹³C values from -19.9 to -10.7‰ (Blanz et al., 2020).

The absence of a consistent corresponding shift in δ^15^N values argues against a simple trophic explanation for the elevated δ^13^C values. While a broad marine diet is typically characterised by enrichment in both δ^13^C and δ^15^N values, the observed δ^15^N values largely overlap with terrestrial livestock baseline values. This decoupling suggests that the δ^13^C enrichment is not a result of a higher trophic position or the unlikely consumption of marine protein, but rather the consumption of a specific ^13^C-enriched plant source with moderate ^15^N values.

### 4.2. Nitrogen variability and flock management

In contrast to the constrained δ^13^C values, δ^15^N values exhibit substantial inter-individual variability both within and between sites. This variability is most clearly expressed at Vester Egesborg and Bejsebakken, which display pronounced intra-site variability, and is also evident at Fredshøj, the site represented by the largest dataset. The presence of both low and elevated δ^15^N values at the same sites indicates that local sheep were exposed to heterogeneous nitrogen isotope baselines, which could include foraging on beach meadows (Prummel et al., 2024; van der Sluis & Reimer, 2021).

It should also be noted that the three individuals with δ^15^N values exceeding 9‰ may represent suckling or recently weaned lambs. Elevated δ^15^N values in juvenile mammals are a well-established trophic effect resulting from the consumption of maternal milk, which is enriched in ^15^N relative to the maternal diet (Schurr, 1998). Zooarchaeological age assessments indicate that two of these individuals, from Vester Egesborg and Fredshøj, were approximately 6-12 months old (Jæger & Gotfredsen, unpublished), supporting the interpretation that their elevated δ^15^N values reflect ontogenetic effects rather than differences in grazing location or foddering practices.

This variability may reflect differences in grazing location, access to manured fields or foddering practices. The coexistence of these signals at individual sites suggests that sheep were not managed under a single, uniform regime but rather experienced a range of management practices that may have varied seasonally or spatially across the local landscape.

### 4.3. Sulfur isotopes and landscape use

Within-site comparisons further highlight the apparent complexities of sheep management practices. At Fredshøj and Bejsebakken, variability in both δ¹⁵N and δ³⁴S values indicates that sheep within a single settlement were not managed homogeneously. This pattern is consistent with the subdivision of local flocks across different grazing areas, the use of multiple foddering strategies, or differential access to manured soils within the same community. It should also be noted that sheep may have been brought to the site prior to slaughter; some isotopic variability may therefore reflect the integration of animals raised in different grazing environments.

In contrast, sites represented by one to three individuals yield isotope values that fall within the broader ranges observed across the assemblages. While these values provide useful contextual information, they should not be overinterpreted as representations of site-level management strategies.

### 4.4. Eelgrass-derived carbon as a plausible dietary contributor

One plausible explanation consistent with the observed isotope pattern is the incorporation of eelgrass (*Zostera sp.*)-derived carbon into sheep diets, either through direct grazing in coastal environments or ecotones, or through supplementary foddering with harvested eelgrass (Alm, 2003; van der Sluis & Reimer, 2021). Eelgrass fits the observed isotopic profiles well, as eelgrass is characterised by substantially enriched δ^13^C values (averaging -9.1 ± 2.2‰) relative to terrestrial C_3_ vegetation because it utilises isotopically heavier bicarbonate and marine dissolved inorganic carbon during photosynthesis (Meadows & Fischer, 2024; Papadimitriou et al., 2005). Consequently, in Danish waters, eelgrass leaves can exhibit δ^13^C values as high as -8‰ during summer (Papadimitriou et al., 2005). This heavy signal is readily transferred through the food web, resulting in consumer values markedly enriched relative to the typical ranges reported for ovicaprines in Scandinavia (Gron et al., 2024; Gron & Rowley-Conwy, 2017; Jørkov et al., 2010; Macheridis et al., 2024; Schjerven et al., 2024; van der Sluis & Reimer, 2021).

Importantly, the dataset shows a decoupling of δ¹³C and δ¹⁵N values, a diagnostic feature of the eelgrass biotope (Meadows & Fischer, 2024). Unlike pelagic food webs, where high carbon enrichment is typically accompanied by increased nitrogen due to longer trophic chains and higher-trophic level prey, the eelgrass food web is relatively short and supported by nitrogen fixed from or absorbed from seabed sediments, which is often depleted in ^15^N compared to open-sea sources (Meadows & Fischer, 2024). Furthermore, while modern seagrass δ^15^N can be anthropogenically elevated by runoff, values in unimpacted waters are significantly lower (Papadimitriou et al., 2005). This is consistent with the terrestrial-like nitrogen values observed in these specimens, reflecting the natural nitrogen baseline of eelgrass meadows.

Sulfur isotope data provide contextual support for this interpretation. While pelagic marine systems reflect uniform oceanic sulfate (approximately +20.3‰), eelgrass roots absorb ^34^S-depleted sulfide generated by sulfate-reducing bacteria in anaerobic seabed sediments (Craig et al., 2006; Meadows & Fischer, 2024; Nehlich, 2015; Richards et al., 2001). This results in a unique “sulfide-derived” signature (ranging from -15 to +10‰) that often overlaps with terrestrial ranges but is distinct from the high values expected from pelagic marine protein or sea-spray effects (Meadows & Fischer, 2024). The combination of elevated δ^13^C and moderate δ^34^S values overlapping terrestrial ranges in the sheep suggests the exploitation of eelgrass.

Distance-to-coast measurements provide additional context for evaluating whether eelgrass consumption reflects direct grazing in coastal environments or the supplementary use of harvested eelgrass. All analysed sites are located within approximately 4 km of the present-day coastline, indicating that coastal environments were broadly accessible across the studied sites. However, the absence of a clear relationship between coastal proximity and δ^34^S values (Figure 6) suggests that eelgrass consumption cannot be explained solely by active grazing in coastal areas. Rather, these patterns are consistent with a model in which eelgrass formed part of an adaptive sheep management strategy incorporating both opportunistic coastal grazing and the supplementary foddering with harvested eelgrass.

**Figure 6.**
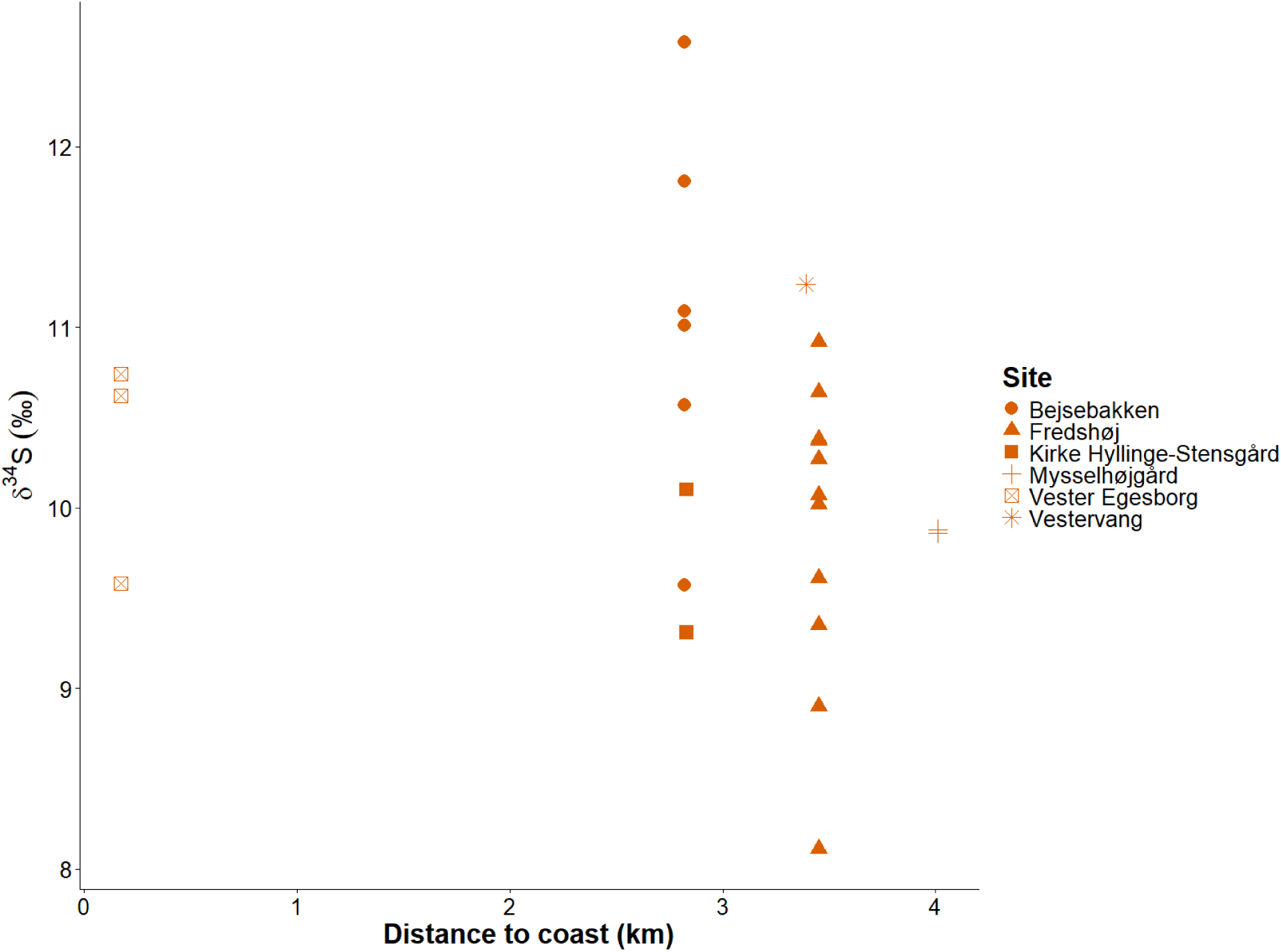
Relationship between distance to nearest coastline and δ³⁴S values in sheep from Late Iron Age Denmark. The lack of a clear relationship between sulfur isotope values and geographic distance from the coast.

**Figure 7.**
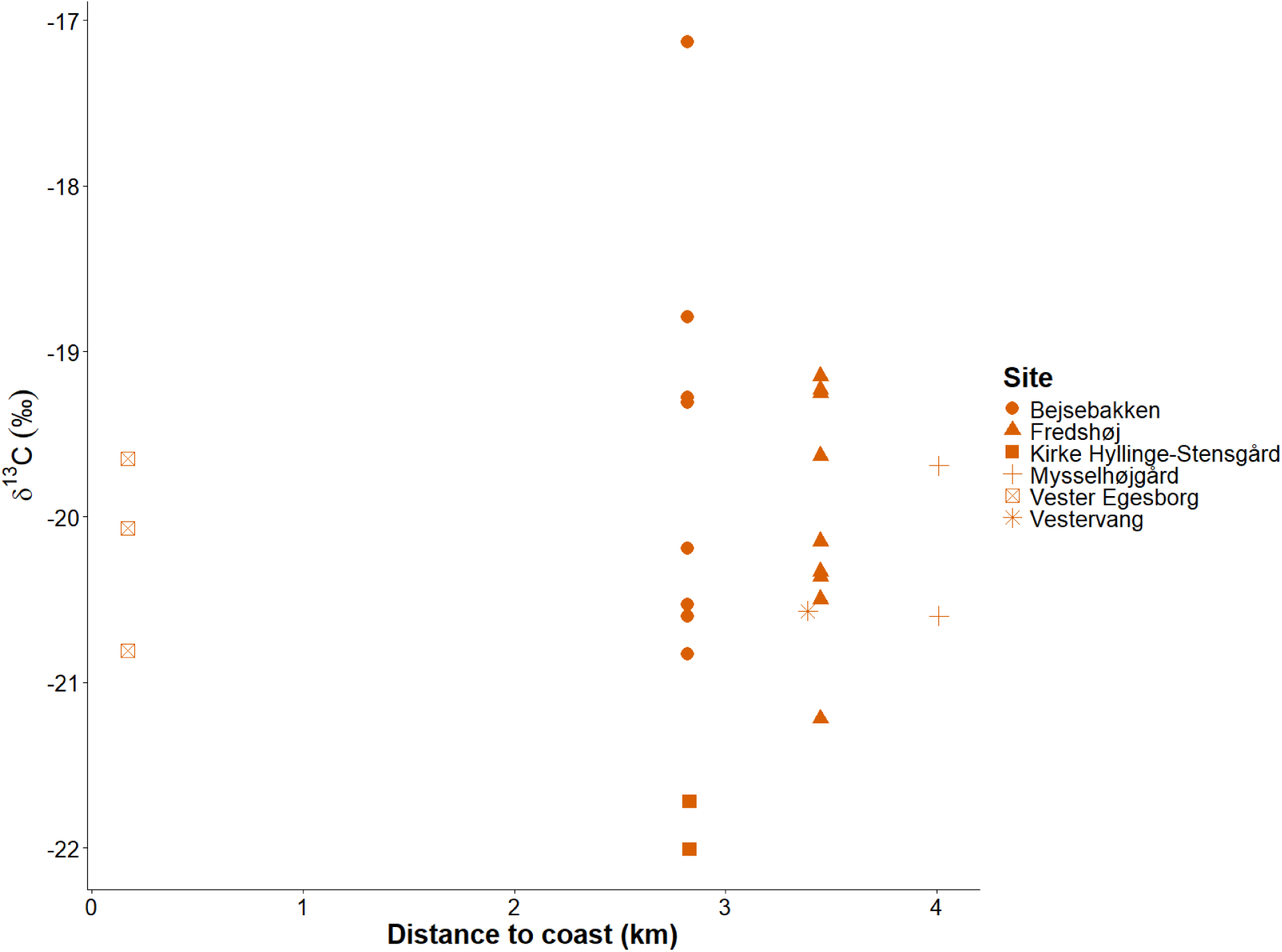
Relationship between distance to nearest coastline and δ¹³C values in sheep from Late Iron Age Denmark. The lack of a clear relationship between carbon isotope values and site-level coastal proximity suggests that elevated δ¹³C values cannot be explained solely by direct grazing in coastal landscapes and is consistent with both opportunistic grazing and supplementary foddering with eelgrass.

A similar absence of a relationship between coastal proximity and δ^13^C values further supports this model, as enriched carbon isotope signals occur across sites regardless of distance from the shoreline.

Given the proximity of all sites to the coastline, freely grazing sheep could have had regular access to coastal grazing areas. Consequently, direct grazing on marine vegetation, including eelgrass, is plausible. The pronounced inter-individual variation observed within sites, combined with the absence of a coastal isotopic gradient, suggests that coastal grazing alone cannot account for the observed isotope patterns. Instead, the isotopic data are more consistent with flexible management strategies in which sheep exploited coastal landscapes opportunistically while also receiving supplementary fodder consisting of harvested eelgrass.

### 4.5. Implications for Late Iron Age sheep husbandry

The sheep husbandry practices implied by the stable isotope data are consistent with archaeological evidence for increasing wool-based textile production during the Late Iron Age, particularly, the Viking Age in Denmark. Textile manufacture during this period intensified both in terms of scale and organisation, as indicated by the increasing number of textile tools and pithouses interpreted as textile workshops (Andersson Strand, 2021). These developments imply not only increased demand for wool but also a need for more resilient and flexible sheep management systems capable of sustaining larger, more productive flocks. Within this context, the apparent integration of coastal plant resources into sheep diets can be seen as part of a broader strategy to stabilise and intensify wool production. Access to eelgrass-rich grazing environments or supplementary fodder derived from coastal vegetation would have allowed sheep to exploit highly productive landscapes that did not compete directly with arable land needed for cereal cultivation.

The archaeological contexts of the analysed assemblages further nuance this picture. The sites included in this study represent a range of settlement types, from ordinary rural settlements to more complex central place environments. This variation is significant, as it provides a framework for understanding how sheep husbandry strategies were embedded within different social and economic settings. While the isotope data reveal broadly comparable patterns of coastal resource use, the organisation, scale, and purpose of livestock management are likely to have differed between these contexts.

The consistency of δ^13^C enrichment across sites suggests that access to ^13^C-enriched coastal vegetation was not incidental but formed a recurring component of sheep management strategies. Rather than reflecting a short-term response to environmental stress, these patterns are more consistent with sustained, structurally embedded practices that supported the maintenance of viable wool-producing flocks. This is particularly relevant given that wool production places different demands on flock composition and longevity than meat-focused husbandry, often requiring the retention of adult animals over multiple years.

At the same time, the pronounced inter-individual variability in δ^15^N and δ^34^S values within individual sites suggests that wool production did not rely on a single, standardised management regime. Instead, sheep appear to have been managed across multiple grazing zones and foddering regimes, potentially reflecting seasonal movement between inland and coastal landscapes or differential household-level access to grazing resources. Such flexibility would have enhanced flock resilience and buffered wool production against local environmental constraints.

When viewed in relation to the settlement context, this variability can be interpreted in different ways. At the rural settlements, isotopic variation is likely to reflect local-scale decision-making, where households balanced access to inland grazing, coastal resources, and foddering strategies in response to environmental and economic constraints. In contrast, sites associated with central place environments, most notably the Lejre complex, including Fredshøj and Mysselhøjgård, may reflect a broader scale of organisation. In such contexts, livestock management was potentially embedded within systems of aggregation and redistribution, where animals were drawn from a wider hinterland and managed in ways that supported more centralised demands, including textile production.

Taken together, the isotope evidence supports a model in which sheep husbandry during the Late Iron Age was integrated with expanding textile economies. Rather than indicating “specialised” flocks fed on marine vegetation, the enriched δ^13^C values reflect the strategic incorporation of coastal plant resources into terrestrial pastoral systems. This flexible exploitation of coastal and inland landscapes would have supported resilient wool-producing flocks during a period when textile production was becoming increasingly central to Scandinavian economies.

## 5. Data availability

All newly generated data are included in the manuscript.

## 6. CRediT authorship contribution statement

**Jonas Holm Jæger:** Conceptualization, Formal analysis, Resources, Data Curation, Writing - Original Draft, Writing - Review & Editing, Visualisation. **Michael P. Richards:** Conceptualization, Investigation, Writing - Review & Editing, Resources, Validation. **Damon C. Tarrant:** Conceptualization, Investigation, Writing - Review & Editing, Validation. **Torben Sarauw:** Writing - Review & Editing, Resources. **Ole Kastholm:** Writing - Review & Editing, Resources. **Julie Nielsen:** Writing - Review & Editing, Resources. **Jens Ulriksen:** Writing - Review & Editing, Resources.

## 9. Acknowledgements

We thank Eva Andersson Strand (Centre for Textile Research, University of Copenhagen), Matthew James Collins and Anne Birgitte Gotfredsen (GLOBE Institute, University of Copenhagen) for their support, discussions and comments on the manuscript.

We also thank Museumsorganisationen ROMU, Museum Sydøstdanmark, Nordjyske Museer, and the Natural History Museum of Denmark for granting access to archaeological material used in this study.

